# Differential Neurodevelopmental Disruption by Bisphenol A (BPA) and Valproic Acid (VPA) in Human Forebrain Organoids

**DOI:** 10.64898/2026.03.15.711882

**Authors:** Mona Zolfaghar, Miaomiao Wang, Lin Li, Moo-Yeal Lee

## Abstract

Neurodevelopmental disorders, including autism spectrum disorder (ASD), are influenced by both genetic abnormalities and environmental toxicants. Among environmental risk factors, endocrine-disrupting chemicals such as bisphenol A (BPA) and pharmaceutical drugs such as valproic acid (VPA) have been associated with an increased risk of autism. In this study, human induced pluripotent stem cell (iPSC)-derived forebrain organoids were used to model early neurodevelopmental disruptions induced by BPA and VPA exposure. On day 62 of differentiation, forebrain organoids were treated with physiologically relevant concentrations of BPA or VPA for 28 days. Following treatment, morphological, molecular, and electrophysiological changes were assessed across experimental conditions. Both compounds produced distinct alterations in organoid morphology, neurodevelopmental gene expression, and network electrical activity, with VPA inducing markedly stronger effects. Overall, these data suggest forebrain organoids as a robust, physiologically relevant *in vitro* model system for studying neurodevelopment. This platform enables systematic investigation of environmental and pharmacological risk factors implicated in the pathogenesis of neurodevelopmental disorders.

## Introduction

Neurodevelopmental disorders, including autism spectrum disorder (ASD), are characterized by impairments in social interaction and communication, along with restricted and repetitive patterns of behavior ^1^. Autism is influenced by both genetic and environmental factors that affect the developing brain ^2^. According to reports from the Centers for Disease Control and Prevention (CDC), autism prevalence in the United States has steadily increased over the past two decades, rising from 1 in 150 children in 2000 to 1 in 36 children in 2020 ^3^. Growing evidence implicates environmental toxicants, including endocrine-disrupting chemicals and pharmacological compounds, as potential contributors to autism pathogenesis. Traditional animal models have provided valuable insights into autism-related neurobiology; however species-specific differences limit their ability to fully capture human-specific developmental pathways and disease phenotypes ^4^. The advancement of human pluripotent stem cell-derived three-dimensional (3D) brain organoid technology offers a transformative platform for modeling early human neurodevelopment *in vitro* ^5^. Increasing evidence highlights the contribution of environmental risk factors to autism pathogenesis. With the availability of human brain organoid models, it is now possible to systematically evaluate how specific chemicals and pharmacological compounds affect early human neurodevelopment. Bisphenol A (BPA) is a widely used industrial compound found in polycarbonate plastics and epoxy resins incorporated into numerous consumer products, including food packaging materials and water bottles ^6^. Due to its extensive use, human exposure to BPA is pervasive, with biomonitoring studies detecting BPA in more than 90% of tested urine samples. This widespread exposure has raised significant public health concerns. Notably, BPA is classified as an endocrine-disrupting chemical. It can interfere with normal hormonal signaling by mimicking, antagonizing, or otherwise altering estrogenic and related endocrine pathways ^7^. The developing brain is especially vulnerable to BPA exposure, as hormonal signaling is essential for regulating neuronal differentiation, synaptogenesis, and overall brain organization ^8^. Disruption of these processes during critical windows of neurodevelopment by BPA can lead to persistent structural, functional, and behavioral abnormalities ^9^. Epidemiological studies have linked prenatal exposure to BPA with an elevated risk of autism in children ^10^. Similarly, valproic acid (VPA) is another well-established risk factor. VPA is a mood-stabilizing medication widely used in the treatment of epilepsy and bipolar disorder ^11^. Pharmacologically, VPA promotes GABAergic neurogenesis and alters epigenetic regulation through inhibition of histone deacetylases (HDACs). HDAC inhibition influences the expression of the transcription factor Pax6, leading to increased expression of glutamatergic proteins ^12^. Prenatal exposure to VPA, particularly during early gestation, has been associated with an increased risk of autism in offspring ^13,14^. Taking VPA during pregnancy, particularly in the second and third trimester, increases the risk of fetal neurodevelopmental disorders by 30 - 40% ^15^. Despite these studies, the molecular, cellular, and functional changes caused by BPA and VPA during neurodevelopment are still not fully understood. In this study, human induced pluripotent stem cell (hiPSC)-derived forebrain organoids were employed to investigate and compare the neurodevelopmental disruptions caused by BPA and VPA. Morphological, molecular, and network-level electrophysiological alterations were systematically analyzed to define compound-specific alterations of autism-associated neurodevelopmental mechanisms. Forebrain organoids were selected for this study because they recapitulate key aspects of the human cerebral cortex, a brain region essential for higher-order cognitive functions, social behavior, and communication, which are commonly affected in autism ^16^. By using forebrain organoids, it is possible to model region-specific neurodevelopmental disruptions and pathway alterations that are highly relevant to the pathophysiology of autism. VPA has been previously studied in brain organoids and shown to induce neurodevelopmental abnormalities consistent with features of autism ^17^. In contrast, the neurodevelopmental effects of BPA have primarily been demonstrated in animal models and two-dimensional (2D) neural cultures, with limited data available in human 3D brain systems ^18–20^. In this study, we employed VPA as a reference compound with well-characterized neurodevelopmental effects to serve as a comparative control for evaluating the impact of BPA on human forebrain organoids. This design enables a direct comparison of two mechanistically distinct neurotoxicants within a physiologically relevant 3D human cortical model.

## Materials and Methods

### Human iPSCs culture

The EDi029A human iPSCs line (Cedar Sinai Biomanufacturing Center), derived from a healthy male donor, was thawed, cultured, and passaged according to the manufacturer’s recommended protocol. Briefly, iPSCs at an initial passage number of 24 were expanded in a 6-well plate pre-coated with 0.5 mg of hESC-qualified Matrigel (Corning, 356234) in complete mTeSR Plus medium (STEMCELL Technologies, 100-0276) and maintained at 37°C in a humidified incubator with 5% CO_2_. The iPSCs were passaged every 4 - 5 days when it reached 70 - 80% confluency, with minimal spontaneous differentiation (<10%), using the StemPro^®^ EZPassage™ disposable stem cell passaging tool (Life Technologies, 23181-010). Cells were passaged as small clumps with a 1:6 split ratio and cultured with daily medium changes.

### Generation of human forebrain organoids from iPSCs

The iPSCs at early passage numbers (up to 32 - 34) stored in a frozen vial were used to generate forebrain organoids according to the published methods with slight modifications ^17,21^. On day 0, iPSCs colonies were harvested using Accutase^®^ (Gibco, A1110501) and resuspended in embryoid body (EB) formation medium consisting of DMEM/F12 (Gibco, 11330032) supplemented with 20% (v/v) knockout serum replacement (Gibco, 10828028), 1x GlutaMAX (Gibco, 35050061), 1x MEM-nonessential amino acids (MEM-NEAA; Gibco, 11140050), 1x Penicillin/Streptomycin (Gibco, 15140122), 2 μM Dorsomorphin (STEMCELL Technologies, 72102), 2 μM A83-01 (STEMCELL Technologies, 72022), 4 ng/mL basic fibroblast growth factor (bFGF; Peprotech, 100-18B), and the CEPT cocktail. The CEPT cocktail consists of 5 μM Emricasan (Fisher Scientific, 50-136-5235), 0.7 μM Trans-ISRIB (Tocris, 5284), 50 nM Chroman 1 (Tocris, 7163), and polyamine supplement diluted at 1:1000 (Sigma-Aldrich, P8483). The collected cells were further dissociated into single cells through pipetting and seeded in an ultralow attachment (ULA) 96-well plate (S-Bio, MS-9096UZ) at a seeding density of 10,000 cells in each well containing 100 µL of the EB formation medium. To maintain the developing EBs, 50 μL of the medium was gently removed from each well and replaced with 50 μL of fresh EB formation medium every 48 hours. The cells were cultured in the EB formation medium supplemented with CEPT and bFGF until day 5, then the medium was replaced with the EB formation medium without CEPT and bFGF for additional 2 days. On day 7, 50 μL of the EB formation medium was removed from each well. Subsequently, 15 μL of 75% Matrigel (Corning, 354230) diluted in DMEM/F12 was added to each well to fully cover the EBs. After 20 minutes of incubation at 37°C to allow Matrigel polymerization, 100 μL of neural induction medium was added to generate neuroectoderm. The neural induction medium consisted of DMEM/F12, 1x N2 supplement (Gibco, 17502048), 1x GlutaMAX, 1x MEM-NEAA, 1x Penicillin/Streptomycin, 1 μM SB-431542 (STEMCELL Technologies, 72232), and 1 μM CHIR-99021 (STEMCELL Technologies, 72052). On day 14, neuroectoderms from the ULA 96-well plate were transferred to a ULA 24-well plate, with each well containing 300 - 500 μL of forebrain organoid differentiation medium to generate forebrain organoids. This medium was composed of DMEM/F12 supplemented with 1x N2, 1x B27 (Gibco, 17504044), 1x GlutaMAX, 1x MEM-NEAA, 1x Penicillin/Streptomycin, and 2.5 μg/mL insulin (Sigma-Aldrich, I9278). The medium was refreshed every 2 - 3 days, and forebrain organoids were differentiated for up to 90 days.

### Immunofluorescence staining of human forebrain organoids

For immunofluorescence staining, forebrain organoids were collected in 1 mL microcentrifuge tubes (ThermoFisher, 05-408-129) and rinsed three times with phosphate-buffered saline (PBS) for 10 minutes each. The organoids were then fixed with 4% paraformaldehyde (PFA; ThermoFisher, J19943.K2) at 4°C overnight. After fixation, they were rinsed three times with PBS for 10 minutes each. Organoids were permeabilized in 0.2% Triton X-100 (Sigma, 9036195) in PBS for 40 minutes at room temperature, followed by incubation in a blocking buffer containing 2% bovine serum albumin (BSA) in PBS for 2 hours at room temperature. Primary antibodies were diluted to their recommended concentrations in PBS containing 0.1% Tween-20 (PBS-T; ThermoFisher, 85113) (**Supplementary Table 1**). The organoids were incubated overnight at 4°C with primary antibodies targeting neural biomarkers, including SOX2 (neural progenitor marker), TBR2 (intermediate progenitor marker), CTIP2 (deep-layer cortical neuron marker), MAP2 (mature neuron marker), FOXG1 (cortical identity marker), PAX6 (cortical progenitor marker), MBP (oligodendrocyte and myelination marker), N-CAD (neural stem cell adhesion and polarity marker), GFAP (astrocyte marker), and TUBB3 (neuronal differentiation marker). Following overnight incubation with primary antibodies, the organoids were rinsed three times with PBS for 10 minutes each and then incubated with the appropriate secondary antibodies diluted in PBS-T for 2 hours at room temperature (**Supplementary Table 2**). After incubation with secondary antibodies, the organoids were washed three times with PBS for 10 minutes each, then incubated with 1 μg/mL DAPI in PBS for 40 minutes to stain nuclei. This was followed by a final wash with PBS three times for 5 minutes each. The organoids were subsequently incubated with a tissue-clearing solution (Visikol Histo-M, HH-10) according to the manufacturer’s instructions. Fluorescence images were acquired using a confocal microscope (LSM710, Zeiss) at 10x and 40x magnifications and analyzed using ImageJ/Fiji v1.54f software.

### Gene expression analysis in human forebrain organoids by RT-qPCR

To confirm the identity of human forebrain organoids, the expression of key neural genes in day 42 forebrain organoids was examined using reverse transcription quantitative PCR (RT-qPCR), with iPSCs serving as the control. For characterization of normal organoid differentiation at day 42, ten organoids were collected. In addition, to investigate the effects of BPA and VPA on brain organoid differentiation, day 90 forebrain organoids exposed to the compounds, along with untreated control organoids, were analyzed. Five compound-exposed organoids from each group were collected. For RT-qPCR analysis, organoids were transferred into 1 mL microcentrifuge tubes and washed three times with PBS. RNA was extracted using the RNeasy Plus Mini Kit (Qiagen, 74134), according to the manufacturer’s instructions. Complementary DNA (cDNA) was synthesized from 1 µg of total RNA using the High-Capacity cDNA Reverse Transcription Kit (Applied Biosystems, 4368814). RT-qPCR was conducted using SYBR™ Green Master Mix (ThermoFisher, A25742) and gene-specific forward and reverse primers (IDT Technologies; **Supplementary Table 3**). Reactions were performed on a QuantStudio™ 5 Real-Time PCR System (Applied Biosystems, A28574) under the following cycling conditions: 40 cycles of denaturation at 95°C for 30 seconds, annealing at 58 - 62°C for 45 seconds, depending on the primer pair, and extension at 72°C for 30 seconds. The expression levels of target genes were normalized to the housekeeping gene glyceraldehyde 3-phosphate dehydrogenase (GAPDH). The panel of analyzed neural genes included CTIP2 (COUP-TF interacting protein 2), FOXG1 (forkhead box G1), MAP2 (microtubule-associated protein 2), PAX6 (paired box 6), SOX2 (SRY-box transcription factor 2), TBR1 (T-box brain protein 1), TUBB3 (tubulin beta 3 class III), SATB2 (special AT-rich sequence-binding protein 2), VGLUT1 (vesicular glutamate transporter 1), SYN1 (synapsin I), NRXN1 (neurexin 1), and GFAP (glial fibrillary acidic protein).

### Assessment of organoid viability

The viability of human forebrain organoids on day 60 was assessed using a fluorescence-based Live/Dead™ viability/cytotoxicity kit (Life Technologies, L3224) containing calcein AM and ethidium homodimer-1 (EthD-1), following the manufacturers’ instructions. Briefly, day-60 forebrain organoids were rinsed with 1x PBS and transferred to a 96-well plate with one organoid per well. Oganoids were incubated with 100 μL of DMEM/F-12 medium containing 2 μM calcein AM and 4 μM EthD-1 for 45 minutes at 37°C in a CO_2_ incubator. DAPI was then added and incubated for an additional 10 minutes. After staining, the organoids were rwashed three times with PBS for 10 minutes each. Fluorescence images were acquired using an automated brightfield and fluorescence microscope (BZ-X800E, Keyence).

### Assessment of neurodevelopmental impact of bisphenol A (BPA) and valproic acid (VPA)

On day 62, human forebrain organoids were treated with 10 μM BPA and 500 μM VPA for 28 consecutive days. These concentrations were selected based on epidemiologically and clinically relevant exposure ranges. BPA concentrations fall within the 1 - 10 μM range reported in human exposure studies, whereas VPA concentrations fall within the 0.5 - 1 mM therapeutic range. A stock solution of BPA (Sigma-Aldrich, 239658) was prepared in DMSO, while VPA (Sigma-Aldrich, P4543) was dissolved in a 1:1 mixture of PBS and DMSO. Control organoids received an equivalent volume of DMSO under the same conditions. Culture media were replenished every 2 - 3 days throughout the treatment period.

### Morphological evaluation of forebrain organoids following VPA and BPA treatment

To evaluate structural changes, forebrain organoids were morphologically assessed after exposure to VPA and BPA on days 0, 7, 14, and 28 and compared with untreated control organoids. Representative brightfield images were captured using a Keyence BZ-X800E microscope. Organoid growth at day 28 was calculated using the following formula: Growth rate (%) = [(X_28_ – X_0_) / X_0_] × 100, where X_0_ denotes the mean organoid size before treatment and X_28_ denotes the mean organoid size at day 28.

### Electrophysiological recording and data acquisition

Extracellular recordings were obtained from human forebrain organoids using a four-wire tungsten electrode bundle (50 µm diameter per wire) connected to an Intan acquisition system. Recordings were conducted in the culture medium under controlled environmental conditions, and the electrode bundle was gently inserted into the organoid tissue to acquire broadband extracellular signals. Signals were sampled at 30 kHz and recorded for 5 minutes per session, yielding broadband segments suitable for downstream single-unit and population analyses. All recordings were performed inside a grounded Faraday cage on a vibration-isolation platform to minimize environmental electrical and mechanical noise. In total, 22 recording epochs were collected from 11 organoids. To estimate baseline system noise, two additional recordings were obtained using the identical experimental configuration in the culture medium without an organoid (“no-organoid” condition).

### Electrophysiological data processing

Raw extracellular recordings (EDF format) were processed in MATLAB using a multi-stage spike-sorting workflow to obtain curated single-unit spike trains. For spike detection, signals were band-pass filtered between 300 - 3500 Hz using a 4^th^-order Butterworth filter to isolate high-frequency components associated with action potentials. Candidate spike events were identified using a two-stage detection procedure that combined a nonlinear energy operator (NEO)-based threshold with a band-pass negative-peak threshold. The NEO detection threshold was set to 5.0 times the robust estimate of background activity, and candidate events were further required to exceed a band-pass negative-peak threshold of 3.5 robust standard deviations. A refractory exclusion window of 1.0 ms was applied to prevent duplicate detections. To improve temporal precision, each detected event was realigned to the most negative band-pass sample within a ± 0.5 ms window around the initial detection time. Waveform snippets spanning 1.0 ms before and 2.0 ms after the aligned spike time were extracted for downstream analysis. Spike sorting was performed independently for each channel using Gaussian mixture model (GMM) clustering. Cluster number was selected by Bayesian information criterion (BIC) over candidate models ranging from K = 1 to 6, with 5 replicate fits per model and covariance regularization set to 10^-6^. Spike waveforms were represented using the first 6 principal components together with waveform shape features, including trough-to-peak duration, half-width, and peak-to-peak amplitude. Cluster assignments with posterior probability below 0.60 were labeled as uncertain. For channels with fewer than 10 detected spikes, clustering was not further subdivided. Candidate units were subsequently curated using quantitative quality criteria, including spike count, refractory-period violations, cluster confidence, and waveform consistency. Template similarity-based merging was additionally applied to reduce potential duplicate units prior to final review. Units passing predefined objective criteria were further examined using standardized diagnostic plots, including waveform overlays, inter-spike interval (ISI) histograms, and autocorrelograms (ACG). Final unit inclusion was determined by the intersection of automated curation and manual validation. Curated units were exported with spike timestamps, template waveforms, and unit-level metadata for downstream statistical analyses.

### Statistical analysis

Statistical analyses were conducted using GraphPad Prism version 9.3.1 (GraphPad Software). All experiments were performed in triplicate, and data are presented as mean ± standard deviation (SD). P-values were determined using unpaired t-tests, one-way or two-way ANOVA, as appropriate. Statistical significance was defined as follows: **** p < 0.0001; *** p < 0.001; ** p < 0.01; * p < 0.05; and ns (not significant) for p > 0.05. Sample sizes (n) for each experiment are provided in the corresponding figure legends.

## Results

### Generation and morphological characterization of forebrain organoids

Upon differentiation, human iPSCs formed uniform embryoid bodies (EBs) in the ULA 96-well plate, which progressed through the expected stages of neural and regional patterning. Brightfield imaging revealed the progressive development of EBs with smooth, well-defined edges by day 7. These EBs subsequently differentiated into neuroectoderms and matured into forebrain organoids. The forebrain organoids exhibited continuous radial expansion and increased structural complexity, with a denser, layered morphology becoming apparent by day 28. Throughout the differentiation timeline, the forebrain organoids maintained their structural integrity (**Figure 1**).

**Figure 1.**
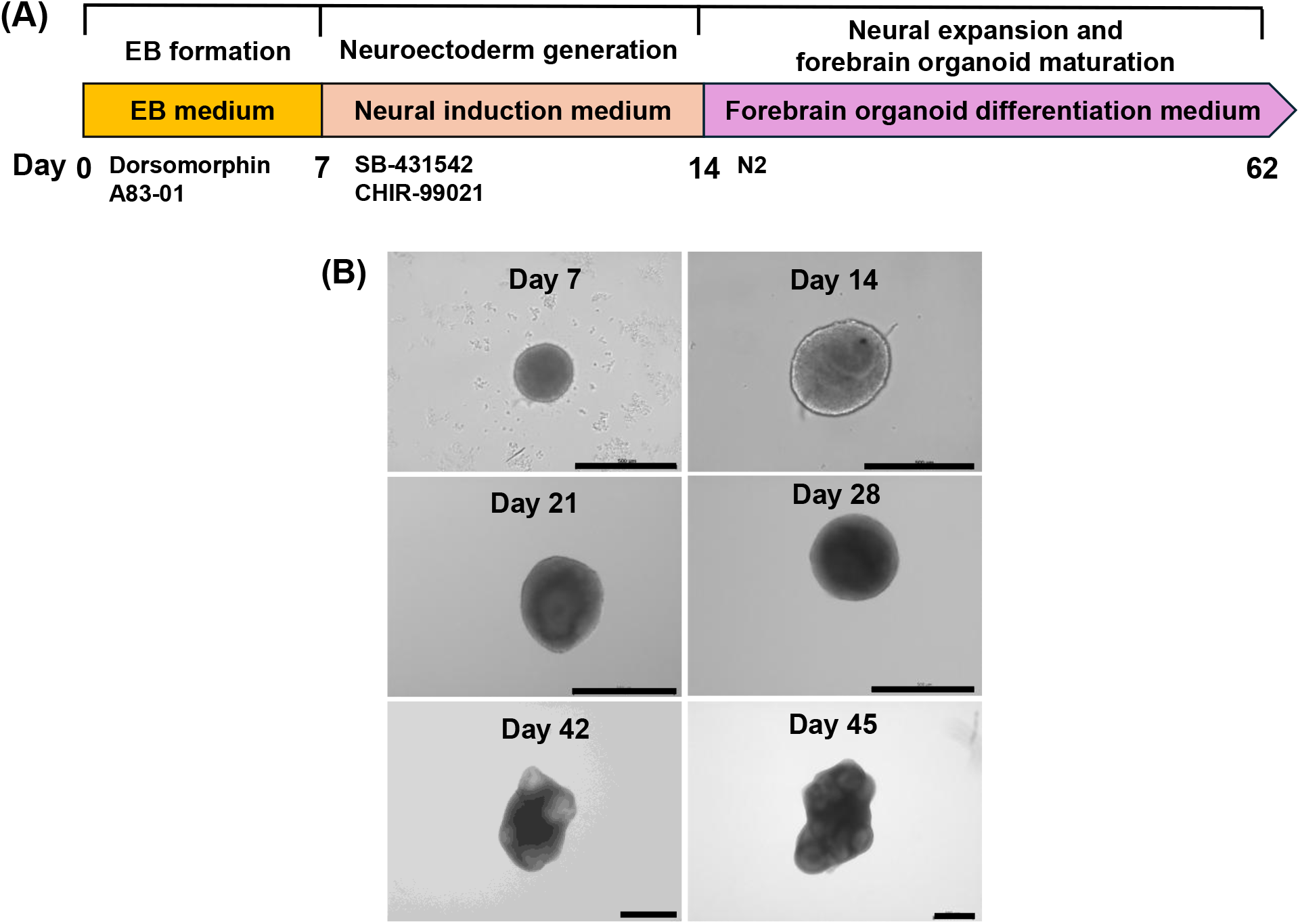
Generation of human forebrain organoids from EDi029A iPSCs. **(A)** Schematic of the differentiation workflow for generating forebrain organoids in an ultralow-attachment (ULA) 96-well plate. **(B)** Representative brightfield images showing morphological progression of human forebrain organoids over time. Scale bars: 500 μm.

### Characterization of forebrain organoids *via* immunofluorescence staining

Immunofluorescence staining was performed on day 42 to confirm the generation of forebrain organoids. Human forebrain organoids were positive for early neural progenitor markers SOX2 and PAX6, indicating the presence of neuroepithelial and radial glial-like cells. The intermediate progenitor marker TBR2 and the deep-layer cortical neuron marker CTIP2 were also detected, confirming progressive cortical neurogenesis. Neuronal maturation was further supported by the expression of MAP2 and TUBB3, which mark dendritic and axonal compartments, respectively. In addition, FOXG1 expression confirmed forebrain regional identity, while GFAP and MBP expression indicated astrocytic and oligodendrocyte lineage commitment, suggesting early glial differentiation. N-cadherin (N-CAD) expression highlighted apical-basal polarity and structural organization. Taken together, these results validate the generation of multicellular brain-like tissue (**Figure 2**).

**Figure 2.**
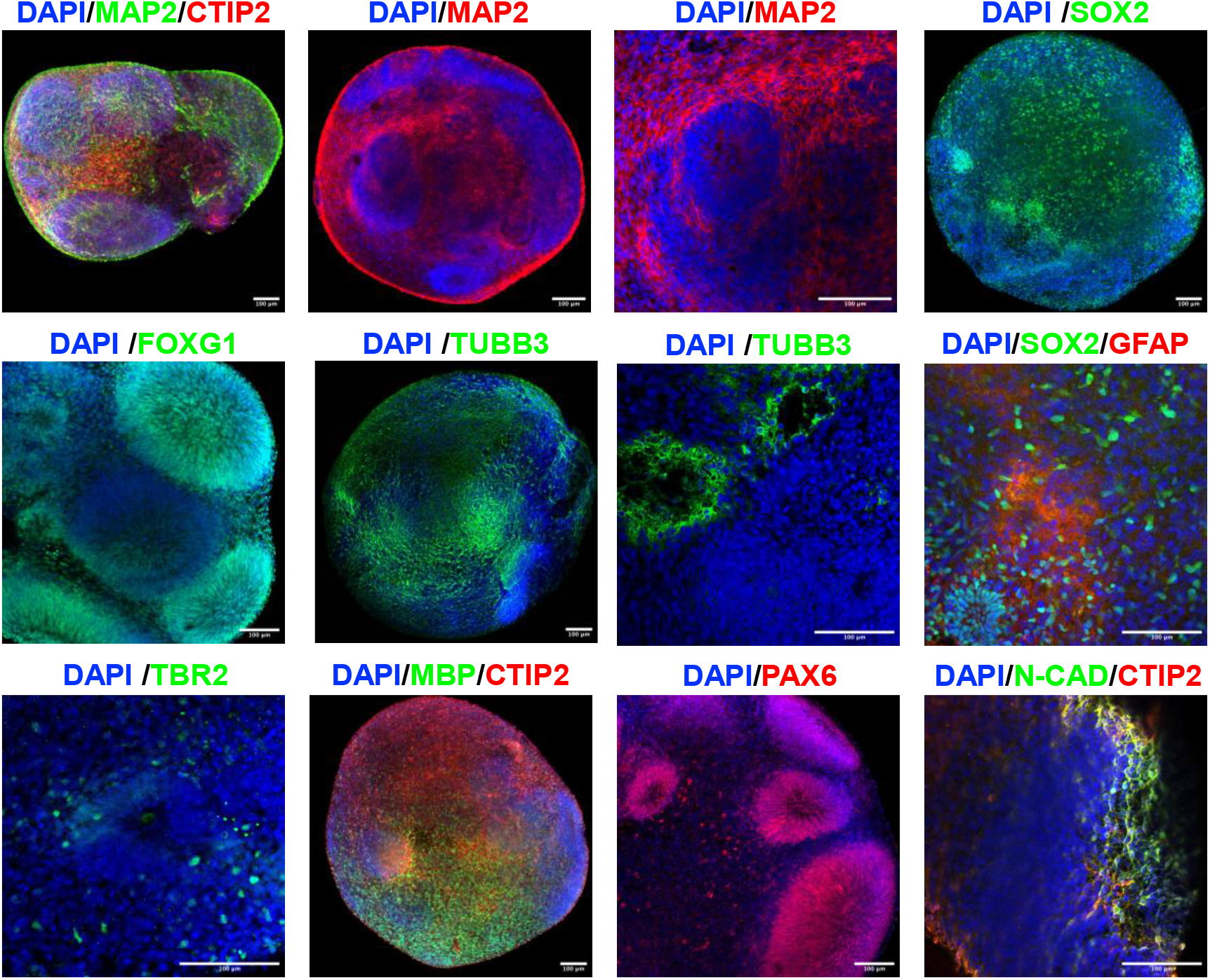
Immunofluorescence characterization of human forebrain organoids at Day 42. Organoids were immunostained to assess neural differentiation, cortical layer specification, and cellular heterogeneity. Markers included MAP2 (mature neurons), TUBB3 (neuronal differentiation), CTIP2 (deep-layer cortical neurons), TBR2 (intermediate progenitors), PAX6 and SOX2 (neural progenitors), FOXG1 (forebrain identity), GFAP (astrocytes), MBP (oligodendrocytes and myelination), and N-cadherin (neuronal cell adhesion). Immunostaining confirmed the presence of diverse neural cell populations and forebrain-specific regionalization within the organoids. Scale bars: 100 μm.

### Validation of forebrain organoid differentiation through RT-qPCR

RT-qPCR analysis performed on day 42 forebrain organoids revealed significantly higher expression of neural genes compared with undifferentiated iPSCs. In particular, *FOXG1, PAX6*, and *TBR1* exhibited strong expression levels. These genes are key regulators of cortical development in the dorsal forebrain. *PAX6*, which is expressed in radial glial progenitors, plays an essential role in maintaining neuroepithelial identity and promoting cortical progenitor expansion. During forebrain development, *PAX6* is expressed in a dorsal-high to ventral-low gradient within the telencephalon, where it regulates genes responsible for dorsal identity and contributes to cortical regionalization ^22^. *FOXG1*, expressed in cortical progenitors that give rise to deep-layer cortical neurons, plays a critical role in telencephalic specification and supports forebrain patterning and cortical growth ^23^. *TBR1*, a marker of early-born deep-layer cortical neurons, showed the highest expression among the analyzed genes and is required for proper cortical lamination and corticothalamic neuron identity ^24^. In addition, genes associated with neuronal differentiation and maturation were strongly expressed. *MAP2* and *TUBB3*, both neuronal cytoskeletal markers, showed elevated expression, indicating neuronal maturation and neurite development. *CTIP2* and *SATB2*, transcription factors marking deep- and upper-layer cortical neurons, respectively, were also significantly expressed, reflecting progressive laminar organization within the organoids. Markers of synaptogenesis and neuronal connectivity, including *SYN1* and *VGLUT1*, were upregulated, suggesting the initiation of functional synaptic network formation by day 42. Although *SOX2* expression remained relatively stable, its persistence indicated the presence of a progenitor population capable of ongoing neurogenesis. Overall, these data highlight the organoids’ fidelity in modeling early human cortical development, with genes such as *TBR1, FOXG1, PAX6*, and *MAP2* serving as key indicators of neuronal identity, cortical patterning, and functional maturation (**Figure 3**).

**Figure 3.**
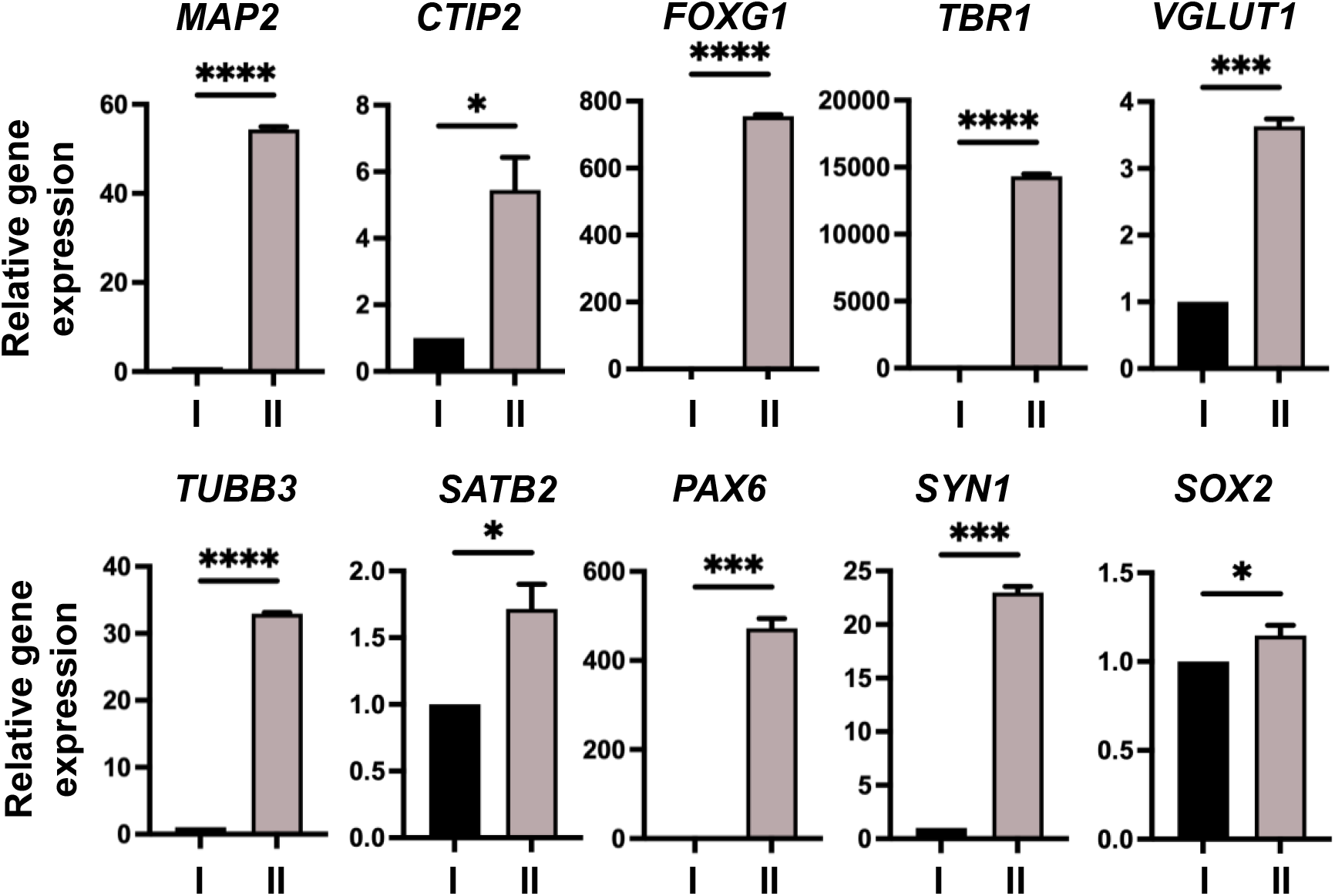
Relative gene expression analysis by qPCR during forebrain organoid differentiation. Relative mRNA expression levels were quantified by qPCR at two developmental stages: **(I)** Day 0 (undifferentiated iPSCs) and **(II)** Day 42 forebrain organoids. Gene expression levels were normalized to housekeeping controls and compared to assess transcriptional changes associated with neural differentiation and forebrain specification.

### Viability of forebrain organoids

Live/dead staining using calcein AM, ethidium homodimer-1, and DAPI in day 60 forebrain organoids demonstrated high cell viability, as evidenced by strong calcein AM fluorescence and minimal ethidium homodimer-1 staining. CellTiter-Glo^®^ 3D assay results further confirmed these findings, indicating robust viability of the organoids at day 60 (**Figure 4**).

**Figure 4.**
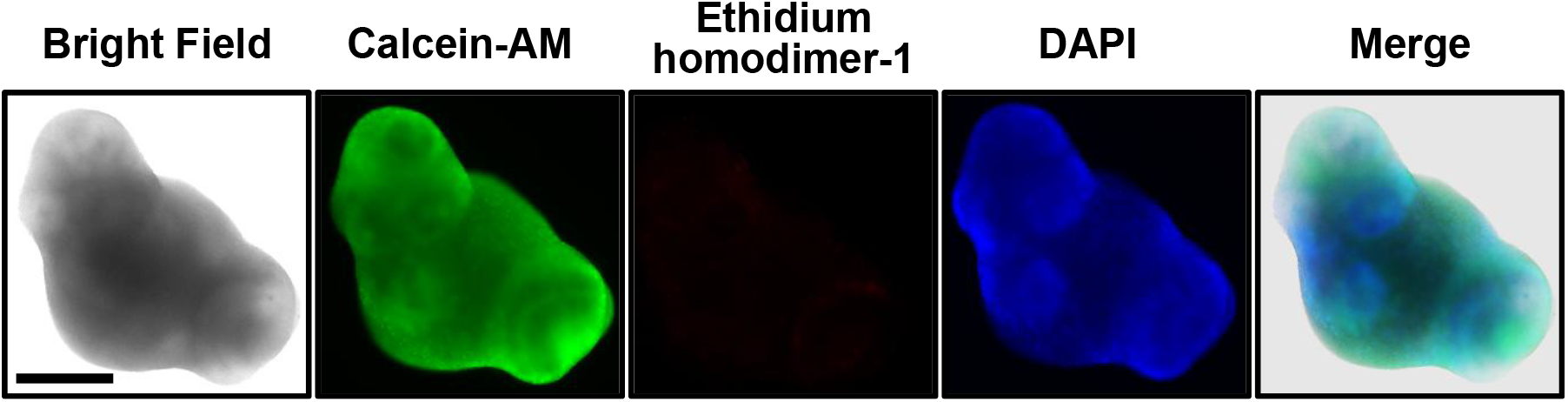
Live/dead viability assessment of forebrain organoids at Day 60. Forebrain organoids were stained with calcein AM to label live cells (green) and ethidium homodimer-1 to label dead cells (red). Fluorescence imaging confirmed high cell viability with minimal cell death. Scale bar: 500 μm.

### Morphological changes by BPA and VPA treatment

Forebrain organoids were treated with 10 μM BPA and 500 μM VPA from day 62 to day 90 of differentiation. Brightfield imaging performed on days 0, 7, 14, and 28 post-treatment revealed treatment-dependent structural changes. Compared with untreated controls, BPA- and especially VPA-treated organoids exhibited reduced size. These morphological differences suggest that prolonged exposure to BPA and VPA may disrupt normal organoid development (**Figure 5A**). Control organoids exhibited robust growth over the 28-day period, reaching a total growth of 43%. In contrast, BPA-treated organoids showed a total growth of 24%, while VPA-treated organoids exhibited a total growth of 20% (**Figure 5B**). BPA-treated and control organoids displayed steady progressive growth throughout the observation period, whereas VPA-treated organoids showed comparatively reduced growth after day 14.

**Figure 5.**
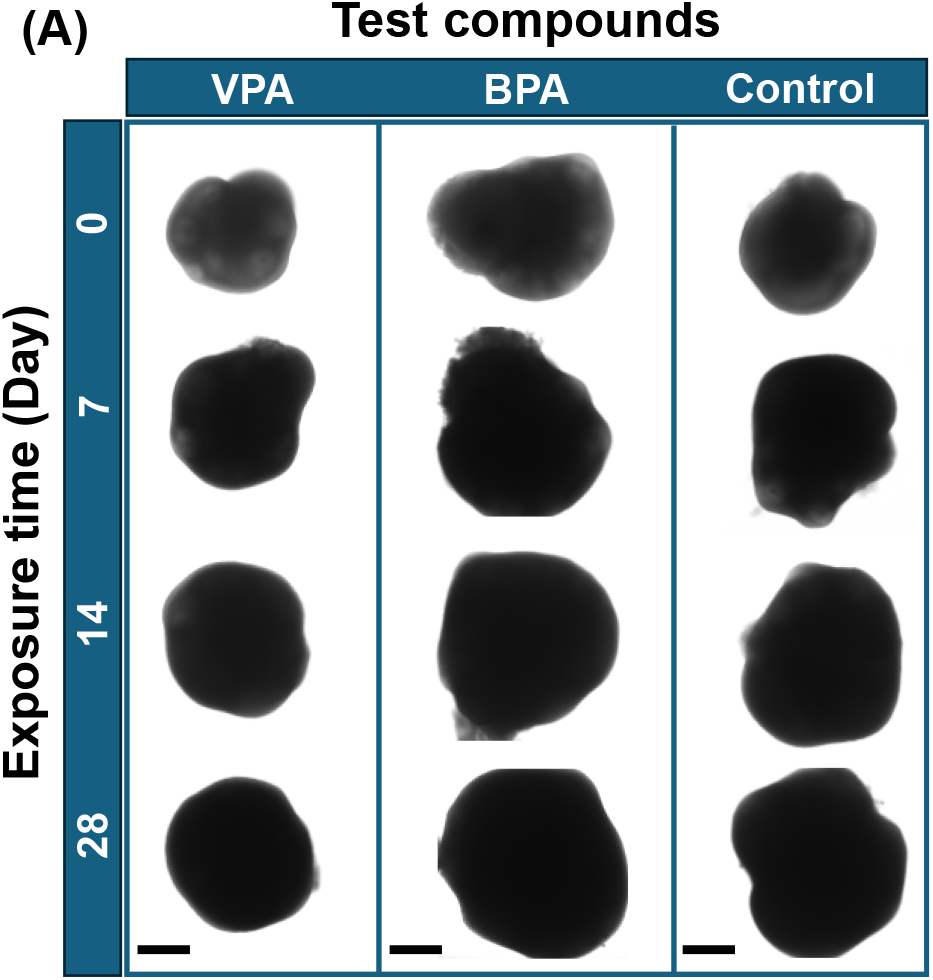
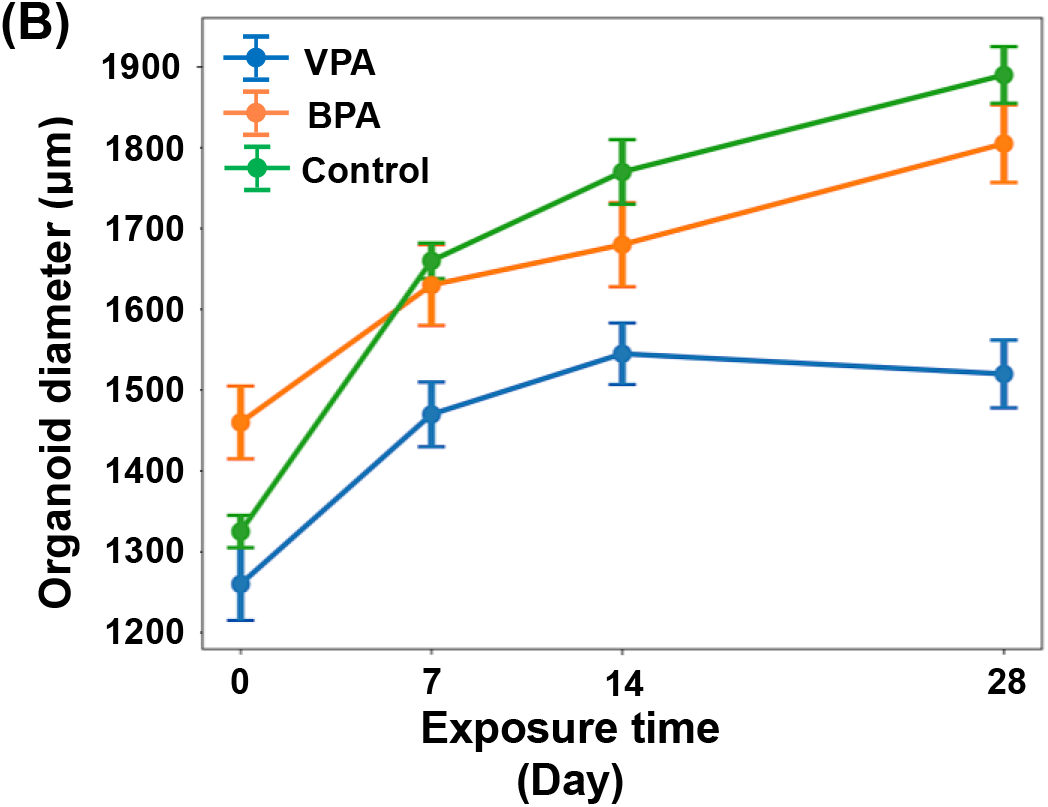
Effects of compound exposure on human forebrain organoid morphology and growth. **(A)** Representative brightfield images showing morphological changes of human forebrain organoids over time in the presence or absence of compound exposure. Treatment was initiated on Day 62 and continued for 28 days. Scale bars: 500 µm. **(B)** Quantification of organoid diameter before and after 28 days of compound treatment, demonstrating changes in growth dynamics. n = 3.

### Gene expression alterations in BPA- and VPA-treated forebrain organoids

To evaluate the effects of VPA and BPA on forebrain organoids and their impact on autism-associated genes, the SFARI Gene database was consulted. Genes classified as high-confidence autism risk genes were selected. Specifically, genes categorized as Score 1 (high confidence), Score S (syndromic), and Score 2S (strong candidate) were included for expression analysis following BPA and VPA exposure. These genes included *TBR1, SYN1, FOXG1, NRXN1*, and *GFAP*, which are key regulators of cortical development and are classified as Score 1. *PAX6*, categorized as Score S, plays an essential role in forebrain patterning and neural progenitor maintenance. *SATB2*, assigned Score 2S, is associated with SATB2-associated syndrome. Although *TUBB3* is not currently scored in the SFARI database, mutations in this gene have been frequently linked to various neurodevelopmental disorders ^25^. To investigate the molecular consequences of prolonged BPA and VPA exposure, RT-qPCR was performed to quantify the expression of neurodevelopmental and autism-related genes in forebrain organoids. Organoids were divided into three groups: untreated control (I), BPA-treated (II), and VPA-treated (III). Significant alterations in gene expression were observed in response to both BPA and VPA exposure, with VPA producing a more pronounced effect. Notably, genes associated with cortical development and neuronal differentiation, including *TBR1, PAX6, TUBB3, FOXG1, NRXN1*, and *SATB2*, were significantly upregulated in the VPA-treated group compared with both control and BPA-treated groups (p < 0.001), suggesting enhanced or dysregulated neuronal specification following VPA exposure.

The synaptic gene *SYN1* was also significantly elevated in the VPA-treated group (p < 0.01 and p < 0.001), indicating potential changes in synapse formation or maturation. *NRXN1* expression increased significantly in the BPA-treated group compared with controls (p < 0.05). *GFAP*, a marker of astrocytic differentiation, was significantly upregulated in the VPA-treated group (p < 0.01), suggesting a potential gliogenic effect of VPA. In addition, *SATB2*, which is involved in cortical neuron identity, showed significant upregulation in both BPA-(p < 0.05) and VPA-treated (p < 0.001) groups, with a stronger effect observed in the VPA-treated organoids. Overall, these results indicate that prolonged exposure to BPA and especially VPA alters the expression of genes involved in neural differentiation, cortical development, and synaptic function (**Figure 6**).

**Figure 6.**
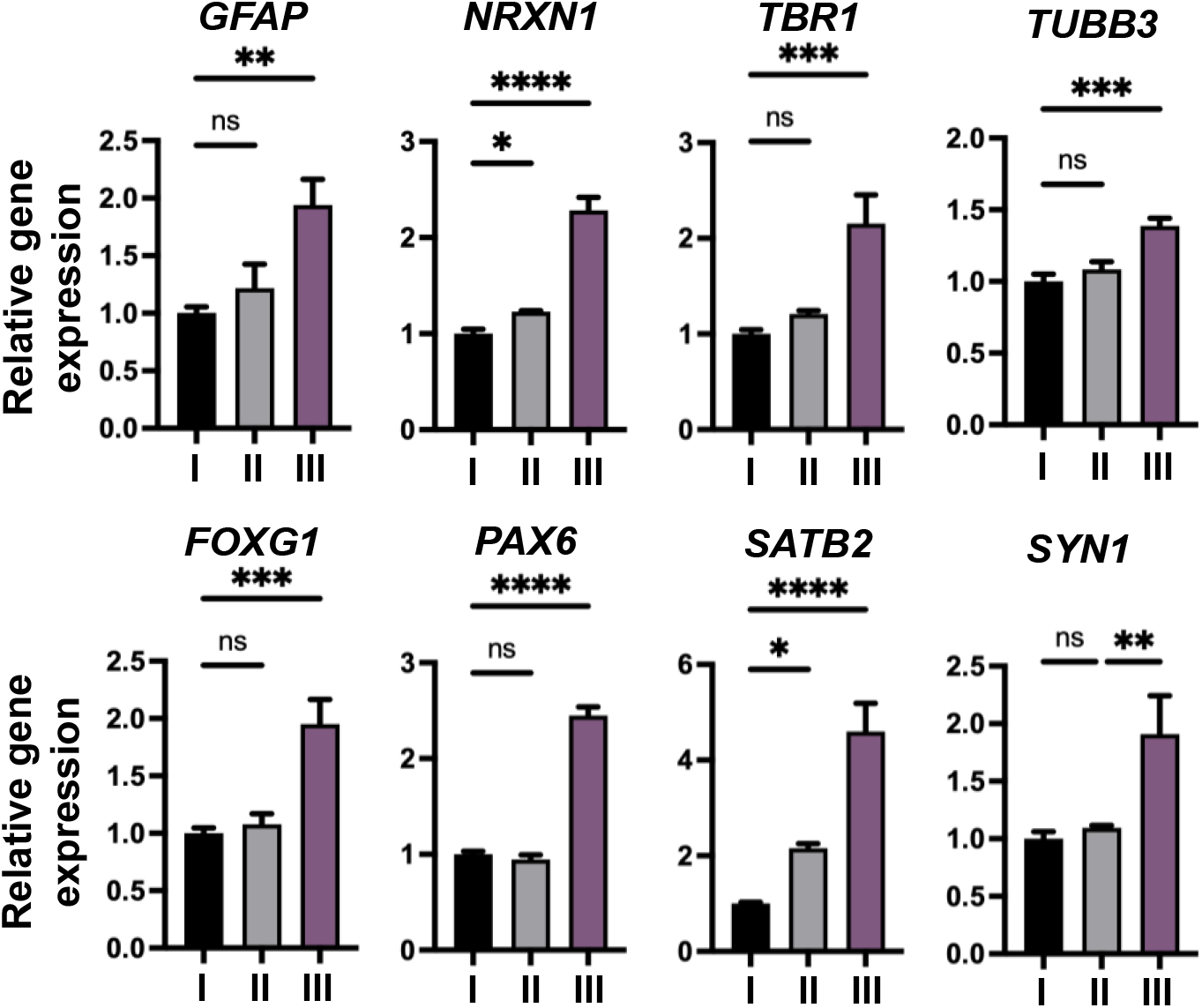
Relative gene expression analysis following compound exposure in Day 62 forebrain organoids. Relative mRNA expression levels were quantified by qPCR in Day 62 forebrain organoids treated for 28 days with (I) no compound (control), (II) BPA, or (III) VPA. Statistical significance is indicated as p < 0.05 (*), p < 0.01 (**), and p < 0.001 (***).

### Electrophysiological alternations induced by BPA and VPA in forebrain organoids

To evaluate how VPA and BPA treatment alters neuronal function in forebrain organoids, probe-based extracellular recordings were performed to monitor spontaneous neural activity. Signals were acquired at 30 kHz, enabling simultaneous analysis of local field potentials (LFPs) and single-unit spiking activity across experimental groups (**Figure 7A**). A total of n = 11 organoids were recorded (Control: n = 4; BPA: n = 4; VPA: n = 3), yielding 39 well-isolated single units for downstream analysis. Representative raster plots and synchronized LFP traces revealed spontaneous firing activity in all groups (**Figure 7B**). Control organoids exhibited temporally structured firing patterns that served as a baseline reference for treatment comparisons.

**Figure 7.**
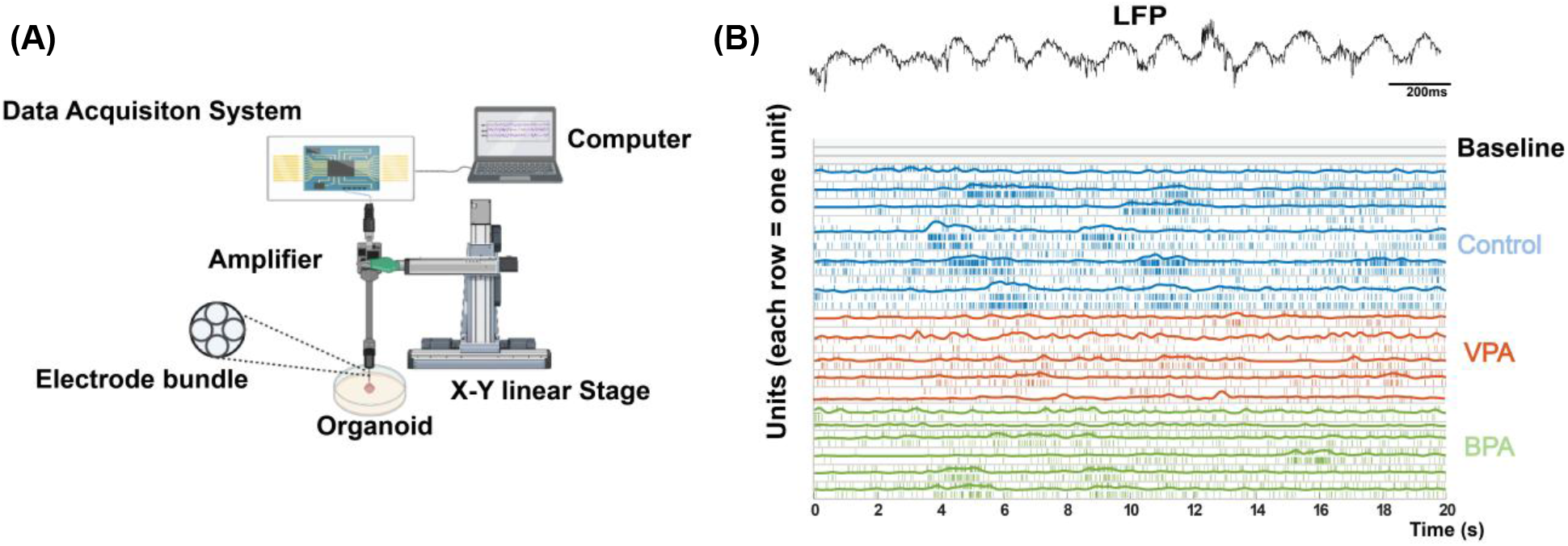
Experimental platform and population-level neural recordings in human forebrain organoids. **(A)** Schematic of the organoid electrophysiology recording setup, showing a four-wire tungsten electrode bundle connected to an Intan acquisition system and recording computer. **(B)** Representative unit activity traces from the baseline (gray), control (blue), VPA (orange), and BPA organoids (green), with synchronized LFP traces shown above. Scale bar: 200 ms.

### Single-unit firing properties

Electrophysiological changes were first examined at the single-unit level. Representative units from each group demonstrated stable spike waveforms, refractory periods in ISI distributions, and well-defined ACG structure (**Figures 8A-C**), consistent with isolation of individual units. Across all detected units (Control: n = 17; BPA: n = 11; VPA: n = 11), mean firing rate differed significantly between groups (one-way ANOVA: F (3, 37) = 7.238, P<0.001). Post hoc comparisons revealed a significant reduction in mean unit firing rate in the VPA group compared with Control (P = 0.006), whereas BPA-treated organoids exhibited an intermediate firing rate not significantly different from Control (P = 0.255) but higher than VPA (P = 0.437) (**Figure 8D**). Because multiple units were recorded from individual organoids, firing rates were additionally summarized at the organoid level to account for potential within-organoid clustering. Unit-level firing rates did not significantly differ across organoids within each group (Control F(3,13)=0.559, P=0.651; VPA F(2,8)=2.707, P=0.127; BPA F(3,7)=1.312, P=0.344) (**Figure 8E**). Kernel density estimates of log10-transformed unit firing rates revealed a leftward shift in the VPA group relative to Control, with BPA showing a partial intermediate shift (**Figure 8F**), indicating redistribution toward lower firing rates rather than complete suppression of activity.

**Figure 8.**
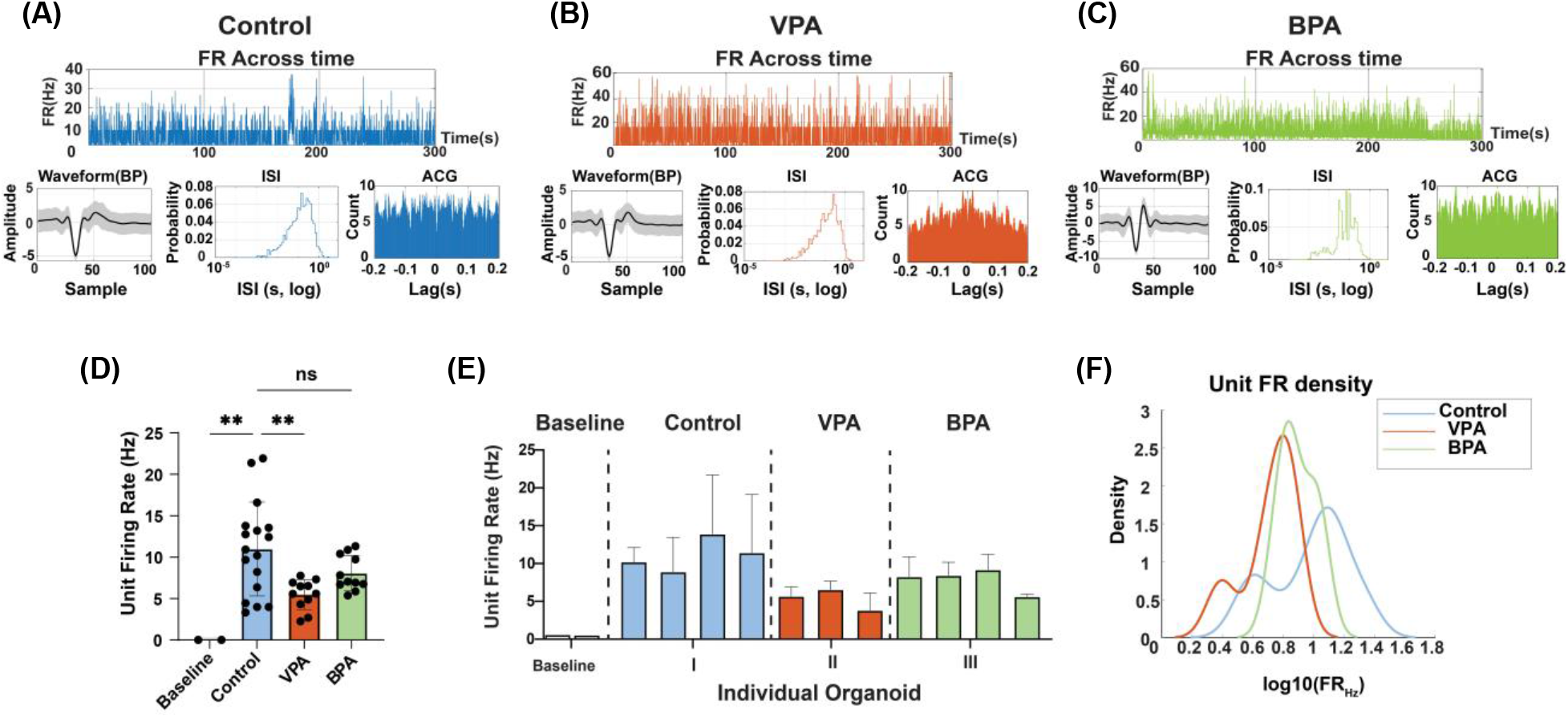
Reduced unit firing rates in VPA-treated organoids and partial preservation in BPA-treated organoids compared with control. **(A-C)** Representative single-unit examples from **(A)** Control, **(B)** VPA, and **(C)** BPA groups. Top: time-resolved firing rate across the recording session. Bottom: mean spike waveform (± s.d.), inter-spike interval (ISI) distribution (log scale), and autocorrelogram (ACG). **(D)** Comparison of mean unit firing rates across groups. Each dot represents one unit (Control: n = 17; BPA: n = 11; VPA: n = 11). Statistical comparisons were performed using one-way ANOVA followed by Tukey’s post hoc test (**P < 0.01; ns, not significant). **(E)** Organoid-level mean firing rate across individual organoids. Bars represent the average firing fate across units detected within each organoid. Dashed lines separate groups. **(F)** Kernel density estimates of log10-transformed unit firing rates for Control, VPA, and BPA groups.

### Population burst dynamics

Network bursting was examined by aligning population firing rate (popFR) traces to burst peaks and comparing burst-centered activity profiles across groups. Control organoids exhibited temporally extended burst profiles characterized by broad peak structures. Burst-aligned heatmaps demonstrated variability across events but preserved overall temporal dispersion, indicating distributed population recruitment (**Figure 9A left**). In contrast, VPA-treated organoids exhibited a temporally compressed burst profile (**Figure 9A middle**). BPA-treated organoids displayed intermediate and heterogeneous burst relative to Control and VPA groups. Quantitative analysis of burst features revealed no significant group differences in burst rate (F (2, 47) = 0.876, P = 0.423) or burst duration (F (2,40) = 0.509, P = 0.605). However, burst width was significantly reduced in both VPA and BPA groups compared with Control (one-way ANOVA: F (2, 46) = 12.74, P < 0.0001; post hoc: Control vs VPA, P< 0.001; Control vs BPA, P

**Figure 9.**
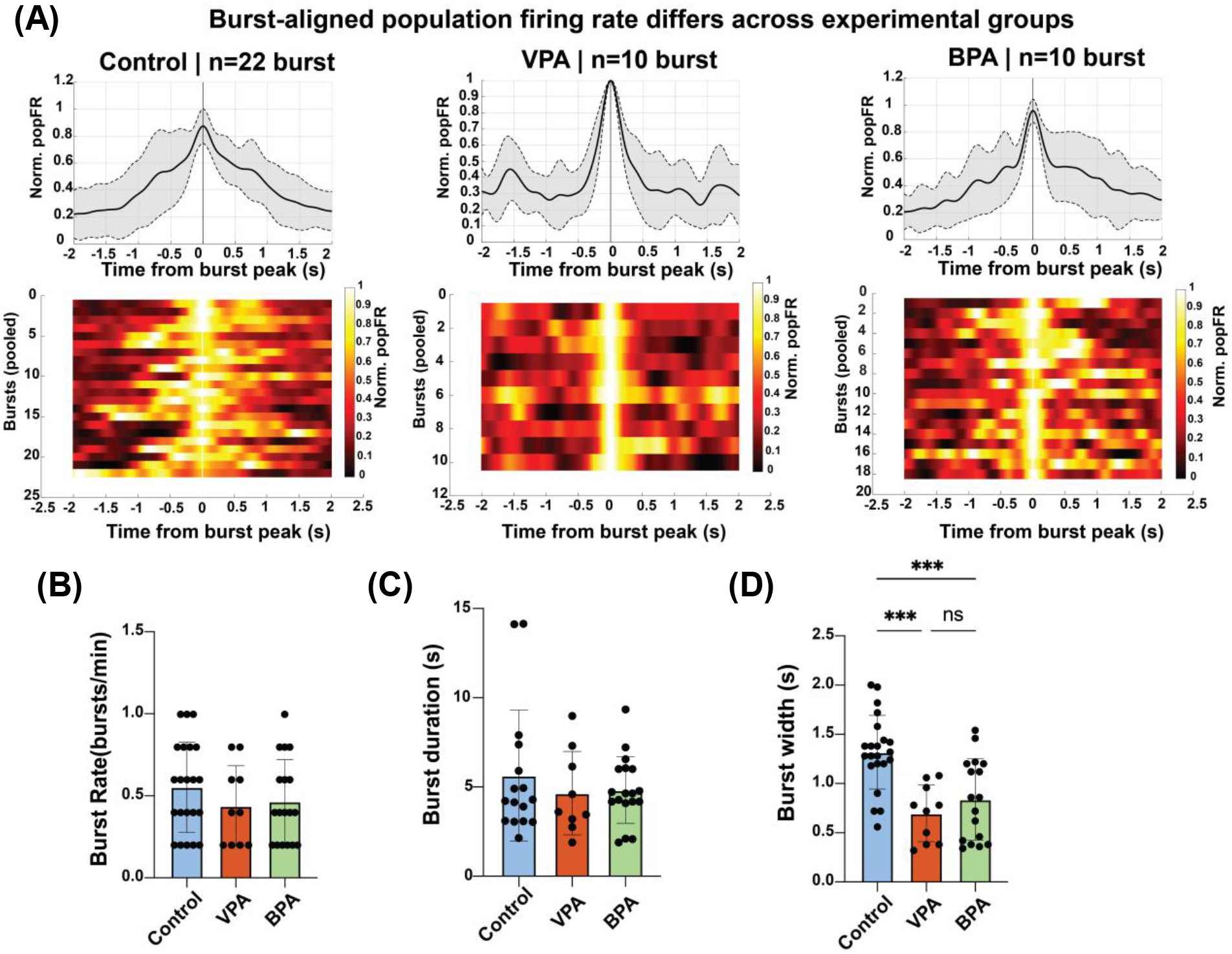
Burst-aligned population firing dynamics differ across experimental groups. **(A)** Burst-aligned population firing rate (popFR) in Control, VPA, and BPA groups. Top: normalized popFR traces aligned to the burst peak (t = 0 s); the solid line indicates the mean across bursts and the shaded area indicates mean ± s.d. across bursts. Bottom: heatmaps showing normalized popFR for each individual burst (rows) as a function of time relative to the burst peak. Color scale indicates normalized popFR (Control: n = 22 bursts; VPA: n = 10 bursts; BPA: n = 10 bursts). **(B-D)** Quantification of burst features across groups: **(B)** burst rate (bursts/min), **(C)** burst duration (s), and **(D)** burst width (s). Each dot represents one burst; bars indicate mean ± s.e.m. Statistical comparisons were performed using one-way ANOVA followed by Tukey’s post hoc test (***P < 0.001; ns, not significant).

## Discussion

Prenatal exposure to BPA and VPA has been associated with an increased risk of autism ^26,27^. However, their direct effects on human brain development remain poorly understood. In this study, forebrain organoids were used to examine the neurodevelopmental impact of BPA and VPA and to compare their levels of developmental neurotoxicity. Unlike traditional 2D cultures or animal models, forebrain organoids offer a 3D, human-specific system. They mimic early brain development and overcome the limitations of previous models, which often lack structural complexity and species specific relevance ^28,29^. Exposure to BPA and VPA from day 62 to day 90 of organoid development resulted in distinct changes in organoid size, gene expression, and electrophysiological activity. BPA- and VPA-treated forebrain organoids exhibited approximately 20 - 24% growth over the treatment period, whereas untreated control organoids demonstrated 43% growth during the same timeframe. Diminished organoid growth is consistent with impaired cortical development and reduced tissue expansion. These phenotypes parallel microcephaly-like features observed in certain neurodevelopmental disorders, including autism ^30,31^. These observations align with previous studies linking environmental stressors to impaired brain growth and disrupted tissue organization ^32,33^. At the molecular level, RT-qPCR analysis of VPA-treated organoids demonstrated significant upregulation of multiple genes strongly implicated in autism according to the SFARI Gene database. Notably, *NRXN1* (Score 1, High Confidence), which encodes a presynaptic adhesion molecule critical for synaptogenesis, has been robustly associated with autism risk ^34^. Dysregulation of key cortical transcription factors was also observed, including *PAX6* (Score S1, Syndromic-High Confidence), a regulator of dorsal forebrain progenitor identity ^35^; *TBR1* (Score 1, High Confidence), essential for deep-layer cortical neuron specification ^36^; and *SATB2* (Score S1, Syndromic-High Confidence), which is expressed in superficial cortical layers and governs upper-layer neuron identity ^37^. In addition, *FOXG1* (Score S1, Syndromic-High Confidence), a master regulator of telencephalic development whose pathogenic variants cause *FOXG1* syndrome with autistic features, was significantly elevated ^38^. Synaptic and cytoskeletal components were similarly affected. *SYN1* (Score 1, High Confidence), involved in synaptic vesicle trafficking and neurotransmitter release, has been linked to epilepsy and autism ^39^, while *TUBB3*, encoding βIII-tubulin required for neuronal differentiation and axon guidance, is essential for proper neurodevelopment ^25^. Furthermore, increased expression of *GFAP* (Score 1, High Confidence), an astrocytic marker, suggests enhanced astroglial maturation or activation. This finding aligns with growing evidence that altered neuron-glia interactions contribute to autism pathophysiology and is consistent with prior reports of VPA-induced astrocytic activation ^40,41^. Compared to VPA, BPA exposure induced comparatively modest transcriptional changes, with significant upregulation limited to *NRXN1* and *SATB2*. Increased expression of *NRXN1*, a synaptic adhesion molecule strongly implicated in autism risk, suggests that BPA may influence early synaptogenic processes ^42^. Similarly, altered expression of *SATB2*, a marker of upper-layer cortical neurons whose disruption has been associated with autism ^43^, points to potential perturbations in cortical projection neuron identity and connectivity ^44^. These results suggest that BPA may influence pathways involved in synapse formation and cortical circuit organization. The attenuated molecular response relative to VPA likely reflects fundamental mechanistic differences between the two compounds. VPA functions as a histone deacetylase (HDAC) inhibitor, broadly modifying chromatin structure and reshaping global gene expression programs. In contrast, BPA does not primarily act through direct epigenetic enzyme inhibition. Rather, it functions predominantly as an endocrine-disrupting chemical, exerting its effects through modulation of estrogen receptor signaling and downstream transcriptional networks ^7^. These distinct modes of action likely account for the more selective and limited gene expression alterations observed following BPA exposure. Additionally, electrophysiological analyses demonstrated that both BPA and VPA altered network activity in forebrain organoids, with VPA producing more pronounced effects on firing rate and burst structure. Control organoids exhibited temporally extended population bursts with broader peak profiles, consistent with previously reported spontaneous network activity in human cortical organoids ^45,46^. VPA exposure was associated with a significant reduction in mean unit firing rate and a temporally compressed burst profile characterized by narrower peak width and faster decay. BPA-treated organoids showed intermediate changes in burst dynamics and firing rate relative to Control and VPA groups. These findings indicate that compound exposure modifies the temporal organization and amplitude distribution of spontaneous network events, with VPA exerting a stronger impact under the present experimental conditions. These observations are broadly consistent with previous studies reporting that VPA exposure can alter excitatory-inhibitory balance and network activity patterns in neurodevelopmental models ^47–49^. BPA induced a milder but significant reduction in burst width, suggesting altered temporal coordination without complete suppression of firing, in line with reports that BPA affects synaptic development and neuronal excitability through endocrine-mediated mechanisms in mice ^50^. Overall, these findings indicate that while both compounds perturb emergent cortical-like network dynamics, VPA exerts a more severe constraint on temporal flexibility and neuronal output. Notably, both VPA and BPA are well-established environmental risk factors for autism and other neurodevelopmental conditions. Our findings reinforce this association. Chronic exposure to these compounds during critical stages of forebrain development induces molecular, morphological, and electrophysiological alterations consistent with disrupted neurodevelopment. These changes mirror key features reported in autism, including altered gene expression, abnormal cortical patterning, and network-level dysfunction. The enhanced expression of genes such as *FOXG1, TBR1*, and *NRXN1*, all of which are associated with monogenic forms of autism, further strengthens the translational relevance of our model ^42,51–53^. By bridging experimental neurotoxicology with human brain organoid modeling, this study emphasizes the urgent need to assess environmental exposures using platforms that more accurately reflect human neurodevelopment. Such systems enable a deeper understanding of how chemical insults during critical periods of brain formation may contribute to the onset of neurodevelopmental disorders, including autism.

## Conclusions

This study demonstrates that prolonged exposure to VPA and BPA induces measurable neurodevelopmental alterations in human forebrain organoids. Both compounds affected organoid growth, gene expression, and neuronal network activity, although VPA produced substantially stronger transcriptional, morphological, and electrophysiological changes than BPA. In particular, VPA exposure was associated with broad alterations in genes involved in cortical development, neuronal differentiation, synaptogenesis, and glial maturation, whereas BPA induced more selective molecular and functional changes. Electrophysiological analyses further revealed that compound exposure alters spontaneous neuronal network activity, with VPA reducing firing rates and compressing burst dynamics, while BPA produced intermediate effects on network organization. These results highlight distinct modes of neurodevelopmental disruption associated with the two compounds. Overall, this work demonstrates the utility of human forebrain organoids as a physiologically relevant platform for investigating chemical effects on early brain development. The integration of morphological, molecular, and functional analyses allows this model to evaluate how environmental and pharmacological factors influence neurodevelopment.

## Supporting information

No link

## Acknowledgements

This study was supported by the National Institutes of Health (R41ES037566 and R16NS131108).

## TOC Graphic

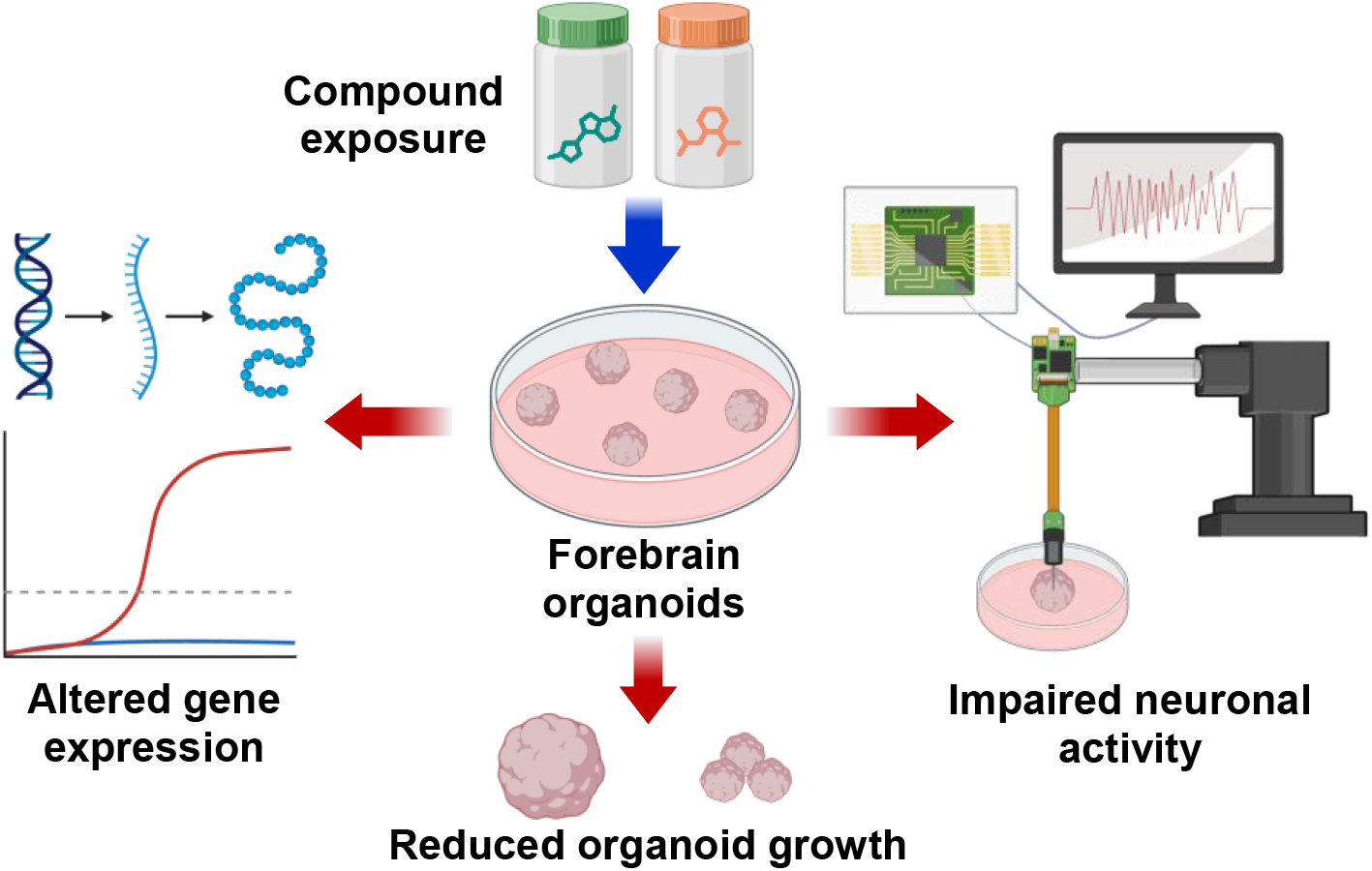

