## Supplementary material for "Differential Neurodevelopmental Disruption by Bisphenol A (BPA) and Valproic Acid (VPA) in Human Forebrain Organoids": No link

### Supplementary Information

**Supplementary Table 1.** List of primary antibodies

| Name | Species | Dilution | Vendor | Catalog # |
|---|---|---|---|---|
| PAX6 | Rabbit | 1:1000 | Abcam | ab195045 |
| SOX2 | Mouse | 1:100 | Santa Cruz | sc-365823 |
| TBR2 | Rabbit | 1:1000 | Abcam | 216870 |
| MAP2 | Rabbit | 1:1000 | Abcam | 183830 |
| CTIP2 | Rabbit | 1:1000 | Abcam | ab 240636 |
| FOXG1 | Rabbit | 1:1000 | Abcam | 196868 |
| N-CAD | Rabbit | 1:1000 | Abcam | ab18203 |
| MBP | Rat | 1:1000 | Abcam | ab7349 |
| GFAP | Rabbit | 1:250 | Abcam | ab 68428 |
| TUBB3 | Rabbit | 1:1000 | Abcam | ab 52623 |

**Supplementary Table 2.** List of secondary antibodies

| Name | Dilution | Vendor | Catalog # |
|---|---|---|---|
| Donkey anti-Rabbit IgG, Alexa Fluor 647 | 1:1000 | ThermoFisher | A-31573 |
| Donkey anti-Mouse IgG, Alexa Fluor 488 | 1:1000 | ThermoFisher | A-21202 |
| Donkey anti-Rabbit IgG, Alexa Fluor 488 | 1:1000 | ThermoFisher | A21206 |
| Donkey anti-Rat IgG, Alexa Fluor 594 | 1:1000 | ThermoFisher | A-21209 |

**Supplementary Table 3.** List of Primers

| Genes | Direction | Primer Sequence (5' - 3') |
|---|---|---|
| *GAPDH* | F | GAAGGTGAAGGTCGGAGTC |
| | R | GAAGATGGTGATGGGATTTC |
| *CTIP2* | F | TCCAGCTACATTTGCACAACA |
| | R | GCTCCAGGTAAGATCGGAAG |
| *FOXG1* | F | CCGCACCCCTCAATGACTTT |
| | R | CCGTCGTAAACTTGCGAAC |
| *MAP2* | F | AGGCTGTAGCACTCCTGAAAG |
| | R | CTTCCTCCACTTGACACTGTG |
| *PAX6* | F | CTGAGGAATCAGGAAAGCAGGC |
| | R | ATGGAGCCCAAGTGGAAGAGG |
| *SOX2* | F | GCTACAGCATGATGCAGGACC |
| | R | TCTGCAGGCTGTCATGAGTT |
| *TBR1* | F | CACTGGAGGTTTGAACGAGG |
| | R | TCTTGGCGCATCCAGTGAC |
| *TUBB3* | F | GGCCAAGGGTCACATCAACG |
| | R | GCAGTCGCAGTTTTTCACATC |
| *SATB2* | F | GAGTGGCAATTCAACCGCAC |
| | R | TCTCGCTCCACTTCTGGGAG |
| *VGLUT1* | F | CAGAGTTTTCGGCTTTCATTG |
| | R | GCACACTGCTTCTAAAGGGC |
| *SYN1* | F | CGATGCCAAATAGTGCAGTG |
| | R | AGCATCGCAGAGGCAGTATTGG |
| *GFAP* | F | CTGGAGGAGAAATTGAGTGC |
| | R | ACGTCAAGCTCCACATGGACC |
| *NRXN1* | F | GCTATCTTGGCAGGTCCTGTGA |
| | R | ACATCCTCAGCCTCCGTATGCA |
